# Neural circuit repair by low-intensity magnetic stimulation requires cryptochrome

**DOI:** 10.1101/424317

**Authors:** Tom Dufor, Stephanie Grehl, Alexander D Tang, Mohamed Doulazmi, Massiré Traoré, Nicolas Debray, Caroline Dubacq, Zhi-De Deng, Jean Mariani, Ann M Lohof, Rachel M Sherrard

**Author notes:** Correspondence to: Rachel M Sherrard, UMR 8256 Biological Adaptation and Ageing, Repairing Neural Networks (R2N), Boite 14, Campus P. and M. Curie, Sorbonne University, 9 Quai St Bernard, 75005, Paris.

## Abstract

Magnetic brain stimulation is a promising treatment in neurology and psychiatry, but clinical outcomes are variable. Unfortunately, mechanisms underlying magnetic stimulation effects are ill-defined, which impedes the development of stimulation protocols appropriate for different neurological conditions. Here we show, *in vivo* and *ex vivo*, that repetitive transcranial magnetic stimulation at low-intensity (LI-rTMS) induces axon outgrowth and synaptogenesis to repair a neural circuit. This repair depends on stimulation pattern, with complex patterns being particularly effective, and its mechanism requires the presence of cryptochrome (Cry), a putative magneto-receptor. Effective LI-rTMS patterns altered expression of Cry target genes known to promote neuronal repair. Because LI-rTMS generates electric fields too weak to depolarise neurons, these findings indicate that the magnetic field itself induces the repair. Our data open a new framework for magnetic stimulation - cryptochrome-mediated molecular and structural neuroplasticity. This information suggests new routes to treatments specific for each neurological disease.

---

Because of our imperfect understanding of the extraordinarily complex human brain, repairing its damage or dysfunction remains one of the major challenges in biomedical science, despite pharmaceutic, stem cell and neuroimaging advances. Non-invasive brain stimulation (NIBS) is increasingly used in neurology and psychiatry in an attempt to trigger intrinsic brain-repair mechanisms. While clinical outcomes are promising, they are variable and underlying mechanisms poorly understood, which hinders therapeutic development^1^. A common NIBS protocol applies magnetic stimulation, which has two forms; (a) repetitive transcranial magnetic stimulation (rTMS) delivering strong magnetic pulses (0.5-2 Tesla, T) to depolarize underlying neurons and trigger activity-dependent plasticity^1^; or (b) low-/pulsed-field magnetic stimulation involving weak pulses (μT-mT) delivered to the whole brain to modulate neuronal function without direct neuronal firing^2–4^. Nevertheless, it is unknown *how* these weak fields modify brain circuits, which are the structural basis of human behaviour.

The biological effects of magnetic stimulation depend on stimulation frequency, pattern, duration and number of pulses delivered^1,5^. These are restricted in high intensity rTMS for technical reasons (heating) and safety considerations (pain, seizures)^6^. In contrast, low field stimulation employs a wide range of frequencies and patterns to modulate neuronal excitability^2,7^, neuroplasticity^8–10^, neuron survival^11,12^, gene expression^10^ and calcium signalling^10,13^.

In this study, we moved beyond individual neurons aiming to identify magnetic stimulation parameters that repair a lesioned neural pathway. For this, we developed focal low-intensity (mT range) repetitive transcranial magnetic stimulation (LI-rTMS), i.e. low intensity magnetic fields delivered focally to only part of the brain like rTMS, while maintaining the safety and frequency range of low-field stimulation^3,6^. Here, we test LI-rTMS protocols for reinnervation in the mouse olivo-cerebellar path and examine mechanisms by which plasticity is induced; information with great clinical significance.

## Results

### LI-rTMS induces olivo-cerebellar reinnervation in vivo

We previously demonstrated that after unilateral olivo-cerebellar lesion (pedunculotomy), intra-cerebellar injection of neurotrophic factors induces remaining inferior olive neurons to grow axon collaterals that partially reinnervate the denervated cerebellar Purkinje cells^14,15^. We have also shown that complex biomimetic high-frequency (BHFS) LI-rTMS prunes abnormal connections in the mouse visual system^8,9^, increases intracellular calcium concentration and modifies neuronal gene expression^10^. We used the same protocol, BHFS 10 min/day for 2 weeks, to stimulate the adult mouse cerebellum after pedunculotomy (Fig 1a&d). Sham stimulation did not induce reinnervation (Fig 1b). BHFS produced VGLUT2-positive terminals in the molecular layer of the left hemicerebellum (Fig 1c) consistent with normal^16^ and BDNF-induced climbing fibres (olivo-cerebellar axon terminals;^14,15^. Labelling extended into the vermis and paravermis and sparsely into the lateral hemisphere (Fig 1e).

**Fig. 1:**
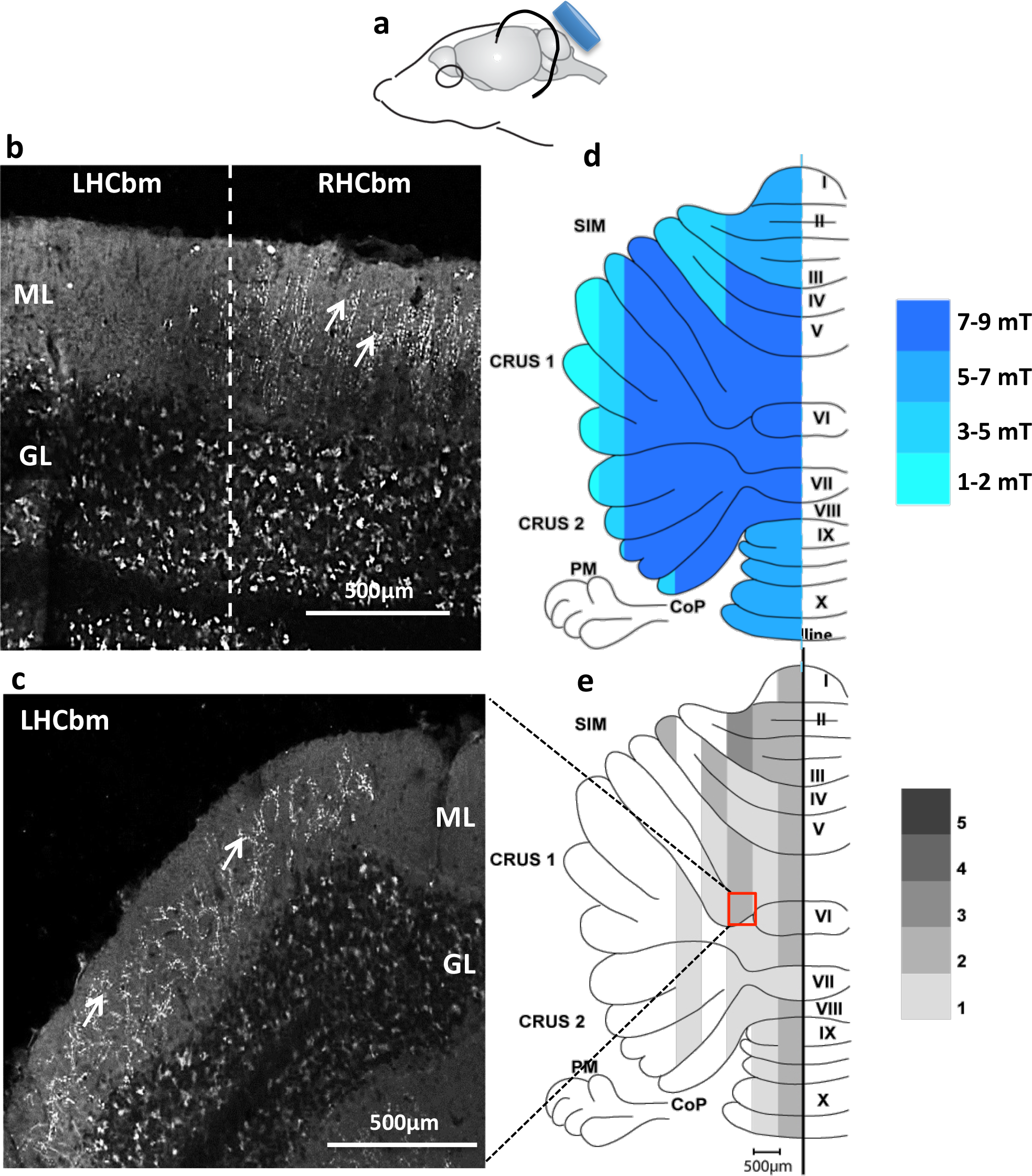
LI-rTMS induces transcommissural climbing fibre reinnervation in adult mice. **a.** Diagram showing the coil (in blue) in relation to the mouse head. **b**, In the cerebellar vermis of a sham-treated pedunculotomized mouse, VGLUT2-positive climbing fibre innervation (thin white vertical lines) is present in the molecular layer of the right hemicerebellum but absent in the denervated left side. Dotted line represents the midline. **c.** VGLUT2-positive reinnervation (thin white vertical lines) is present in the molecular layer of the lesioned left hemicerebellum (lobules simplex) following LI-rTMS. **d.** Diagram of an unfolded cerebellum showing the distribution of magnetic field intensity delivered by LI-rTMS, as measured by Hall device. **e.** The distribution, in 0.5 mm parasagittal zones, of LI-rTMS induced climbing fibre reinnervation. GL = granular layer; ML = molecular layer; LHCbm = left hemicerebellum; RHCbm = right hemicerebellum; SIM = Lobulus simplex; PM = paramedian lobule; CoP = copula pyramidis; I – X = lobules 1-10 of the vermis.

### LI-rMS induces Purkinje cell reinnervation ex vivo in a pattern-dependent manner

Climbing fibre reinnervation *in vivo* (Fig 1) indicates that BHFS LI-rTMS can promote axonal outgrowth. Because reinnervation was incomplete, we tested other LI-rTMS patterns for greater effect. To accelerate the screening of different low-intensity stimulation patterns (LI-rMS, as it is not "transcranial"), we used our *ex vivo* model of olivocerebellar pathway pedunculotomy, denervated-co-cultured hindbrain explants (Fig 2a, Fig S1), in which VGLUT2-positive terminals localized around the Purkinje cell soma and primary dendrites indicate reinnervation^17^ Fig 2b, Fig S1).

**Fig. 2:**
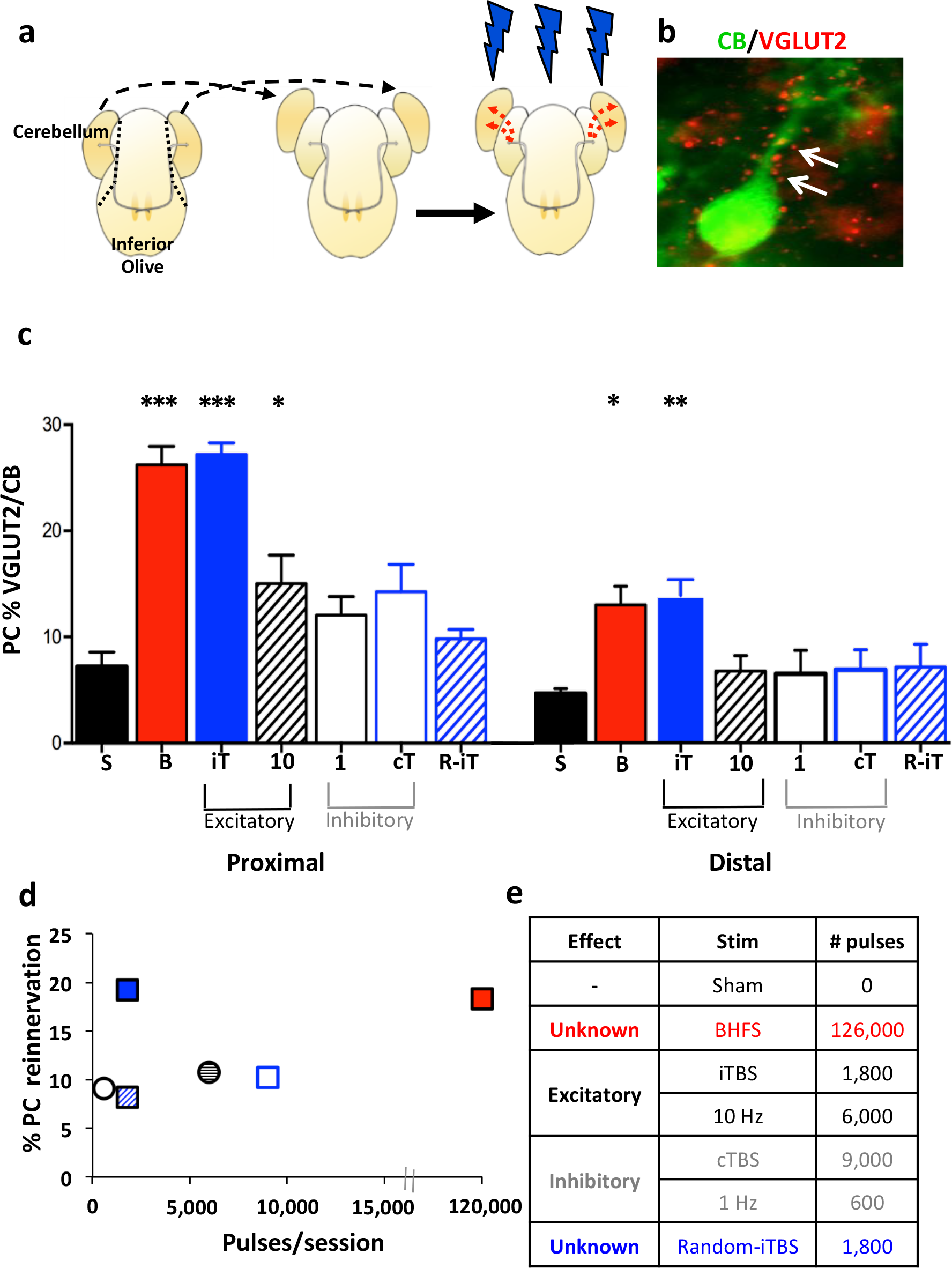
LI-rMS pattern regulates climbing fibre reinnervation *ex vivo*. **a.** Hemicerebellar plates are removed from an explant (dotted line) and placed next to an intact explant (dashed arrows) for reinnervation (red dotted arrows) by host climbing fibre axons (thin grey arrows). 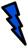. = LI-rMS (see Fig S1). **b.** Purkinje cell (green) showing climbing fibre reinnervation (small red puncta, arrows). **c.** Purkinje cell reinnervation is greater in proximal vs. distal zones of the cerebellar plate (2-way RMANOVA; p=0.000). BHFS (B; n=11) and iTBS (iT; n=8) induced significant reinnervation in both zones compared to sham (S; n=10; ANOVA with Tukey *post hoc*, proximal: BHFS and iTBS both p=0.000; distal: BHFS p=0.003, iTBS p=0.002). 10Hz (n=8) also induced Purkinje cell reinnervation in the proximal zone (p=0.048), which was less than iTBS and BHFS (p=0.001), but not distally (p=0.96). 1Hz (n=6), cTBS (n=8) and random-iTBS (n=7) did not induce significant reinnervation (Proximal zone, compared to sham: 1 Hz, p=0.577; cTBS, p=0.097; random-iTBS, 0.952. Distal zone, compared to sham: 1 Hz, p=0.98; cTBS, p=0.95; random-iTBS, 0.93). Bars are mean  SEM. Significant differences from sham *p<0.05; **p<0.01; ***p<0.001. **d.** Reinnervation density does not correlate with the number of pulses delivered per 10-min session (Pearson coefficient, p=0.353). **e.** Table showing the number of pulses delivered in 10 min for each stimulation parameter studied, and their effects in high-intensity human rTMS. 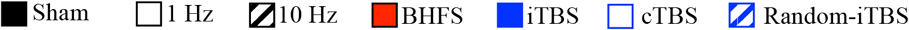

As *in vivo*, BHFS induced reinnervation *ex vivo* (Fig 2c). We then tested frequencies used in human rTMS for facilitation (10Hz and intermittent theta-burst stimulation = iTBS) or inhibition (1Hz and continuous theta-burst = cTBS) of cortical excitability^1^. In each group, VGLUT2 labelling was greater near the host explant (proximal zone) and decreased with distance from the host-graft interface (Fig 2c); inter-group differences persisted in both proximal and distal zones. LI-rMS with iTBS significantly increased reinnervation (Fig 2c), similar to BHFS, but 10Hz was less effective, inducing reinnervation only to the proximal zone (Fig 2c). In contrast, 1Hz and cTBS did not induce reinnervation more than sham controls (Fig 2c).

Next we tested whether the efficacy of complex pattern LI-rMS depended on the number of pulses delivered and found that reinnervation density did not correlate with the number of stimulation pulses (Pearson coefficient, p=0.353; Fig 2d&e). We confirmed this using randomized iTBS (random-iTBS), which delivers the same number of high-frequency bursts in the 2s stimulation, but repeats them randomly (2-60 Hz) rather than at the theta rhythm (5Hz). Two weeks’ random-iTBS failed to induce reinnervation (Fig 2c).

We also examined the target of stimulation: did reinnervation require stimulation of both the cerebellum *and* the inferior olive, or only one or the other? This question is clinically important because stimulation of a whole system is not always feasible, for example the motor cortex *and* the spinal cord. To address this issue, we shielded either the cerebellar or brainstem portion of the explant with mu-metal (see Methods) during daily BHFS LI-rMS. Neither protocol induced significant reinnervation (Fig 3).

**Fig. 3:**
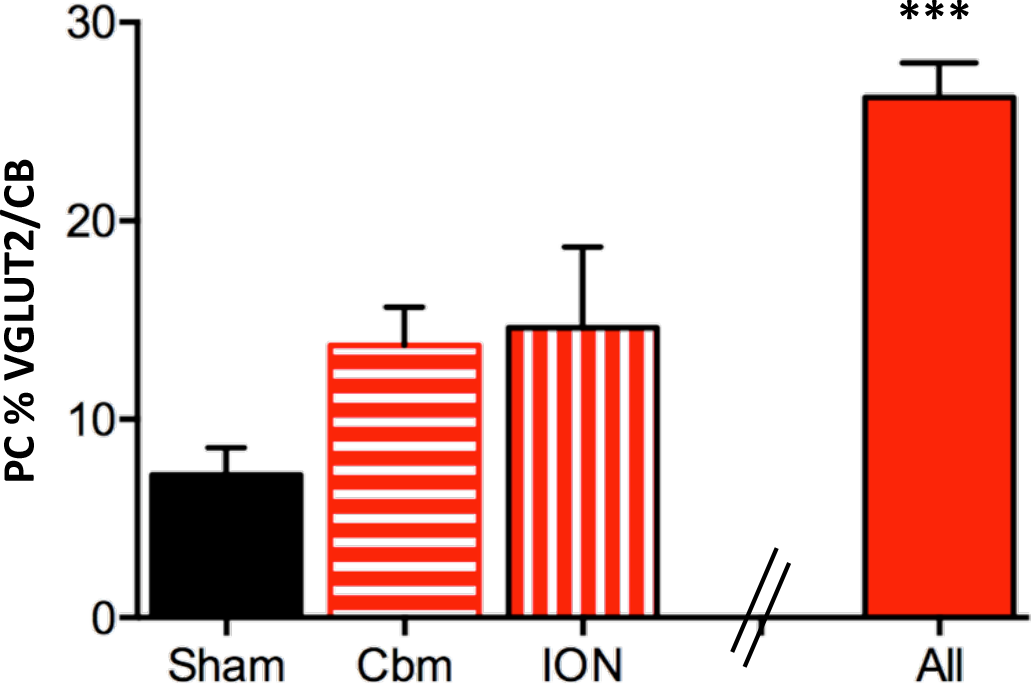
LI-rMS induced reinnervation requires stimulation of both the ION and the cerebellum. BHFS stimulation of either the rostral (cerebellum, Cbm, n=4), or the caudal (ION, n=4) part of the explant does not induce significant reinnervation compared to sham (ANOVA with Tukey *post hoc*; Cbm: p=0.20; ION: p=0.12). “All” is for comparison; showing reinnervation induced by BHFS (***p<0.001) to the whole explant (Fig 2).

Taken together, these studies show that LI-rMS can induce axonal outgrowth and olivocerebellar reinnervation in a frequency-dependent manner. The data also suggest that reinnervation requires stimulation of the whole circuit.

### LI-rMS modulates growth-promoting genes in the cerebellum

To identify potential mechanisms by which LI-rMS brings about neural repair, we examined cerebellar protein and gene expression.

First we used immunolabelling of the immediate-early gene c-fos to identify population(s) of cerebellar cells responding to LI-rMS. A single session of BHFS or random-iTBS doubled the number of c-fos-positive cells (821.3 ±79.7 and 980.8 ± 110 per mm^2^ respectively) compared to sham (439.2 ± 43.1 per mm^2^; p<0.001), whereas iTBS produced only an intermediate increase (599.6 ±64.1 per mm^2^; p>0.05; Fig 4a&b). Thus, the number of pulses given and the percentage of cells expressing c-fos did not correlate (Pearson coefficient, p=0.371; Fig 4c). However, considering only the target cells of climbing fibre reinnervation, Purkinje cells and GABAergic interneurons, BHFS and iTBS *similarly* increased the fraction of cells that was c-fos-positive (Fig 4a&d), whereas random-iTBS did not, confirming the importance of bio-related stimulation patterns.

**Fig. 4:**
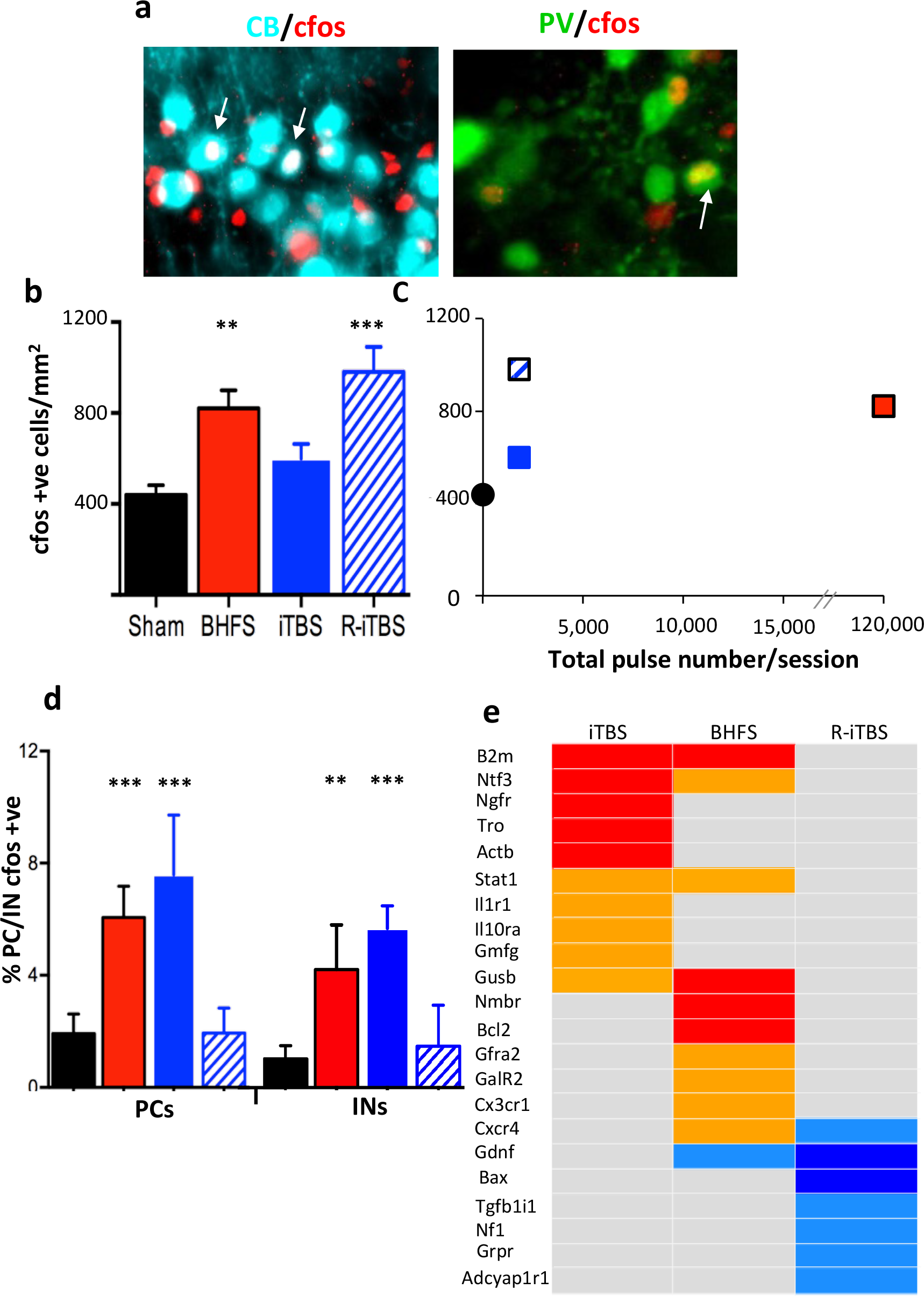
LI-rMS modulates gene expression in denervated hemicerebellar plates. **a.** C-fos (red) in Purkinje cell (cyan) and GABAergic interneuron (green) nuclei (arrows). Bar = 20 μm **b.** The number of c-fos positive profiles per mm^2^ (mean ± S.E.M.) in sham (non-stimulated, n = 8) and BHFS (*n* = 6), iTBS (*n* = 9) or random-iTBS (*n* = 4) stimulated explants. (ANOVA and *post hoc* Tukey. Compared to sham: BHFS, p= 0.001; iTBS, p= 0.16; riTBS, p=0.000). **c.** Stimulation pulse number does not correlate with c-fos labelling (Pearson coefficient, p=0.371). Colour codes as (b). **d.** Percentage of calbindin- or parvalbumin-positive cellular profiles that are double-labelled c-fos/calbindin (“PCs”) or c-fos/parvalbumin (“INs”; mean ± S.E.M.) in sham (non-stimulated) controls or after BHFS, iTBS or random-iTBS. (Fisher’s exact test. BHFS compared to sham: PCs, p=0.000; INs, p=0.006. iTBS compared to sham: PCs, p=0.000; INs, p=0.001. random-iTBS compared to sham: PCs, p=0.827; INs, p=0.682) **e.** Heat-map showing expression changes of 22 genes from the Mouse Neurotrophins & Receptors PCR array after BHFS, iTBS or random-iTBS. Red, upregulated (p<0.05); orange, strong trend for upregulation (0.05<p<0.1); royal blue, down-regulated (p<0.05); light blue, strong tend for down regulation (0.05<p<0.1); and grey, no change. CB = calbindin; IN = interneuron; PC = Purkinje cell; PV = parvalbumin; R-iTBS = random-iTBS. Compared to sham \*\**p* < 0.01; \*\*\**p* < 0.001 Colour codes as (b).

We then examined the expression of genes associated with neurotrophin signalling, because neurotrophins can induce olivocerebellar reinnervation^14,15^. We isolated RNA from the grafted cerebellum during the process of reinnervation (i.e. 6h after the last of 3 LI-rMS sessions). We found significant changes in the expression of 10 genes (Fig 4e), and likely changes in 12 other mRNAs. LI-rMS frequencies that induced significant reinnervation (iTBS and BHFS) increased the expression of genes involved in Gene Ontology (GO) functions such as “axonogenesis, response to axon injury”. In contrast, random-iTBS either did not change or reduced expression of the examined genes (Fig 4e; Fig S2; Table S1). The products of three genes, whose expression was increased by iTBS and/or BHFS (Fig 4e), promote post-lesion neuronal reinnervation in other neural systems: β-2-microglobulin (*b2m*), β-glucuronidase (*gusb*) and Neurotrophin-3/NT3 (*ntf3*)^18–20^

### Cryptochrome is required for LI-rMS-induced post-lesion repair

LI-rMS affects both neuroplasticity and gene expression despite inducing an electric field sub-threshold for cerebellar neuron firing^21^. To determine whether LI-rMS effects depend on the magnetic field *per se*, we compared its effects in wild-type and homozygous double cryptochrome knockout (*Cry1^-/-^Cry2^-/-^*) explants. Cryptochromes mediate magnetic field dependent behaviours^22^ and neuronal excitation^23^ in *Drosophila*, but are unproven magneto-receptors in mammals.

In denervated-co-cultured explants from *Cry1^-/-^Cry2^-/-^* double knockout mice, 2 weeks daily BHFS failed to induce reinnervation (Fig 5). In contrast, cryptochrome double-knockout explants responded to BDNF with significant reinnervation (Fig 5). Thus cryptochrome magneto-receptors are *required* for LI-rMS-induced plasticity but not for plasticity induced by BDNF.

**Fig. 5:**
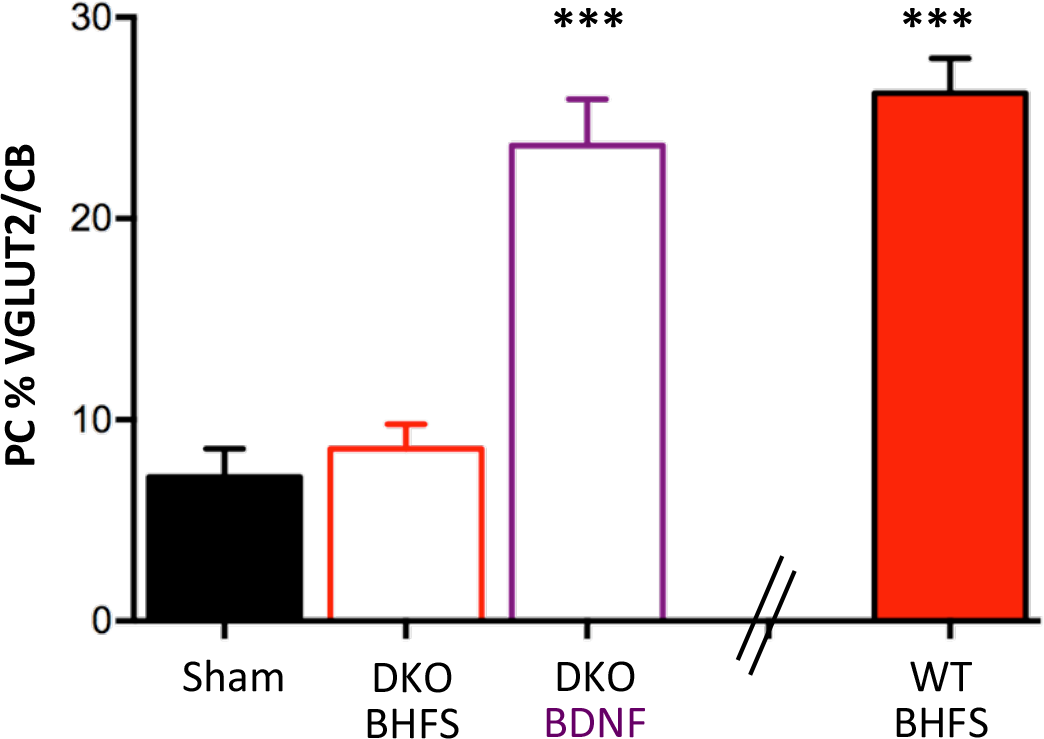
Cryptochromes are required for LI-rMS induced post-lesion repair. Percentage of reinnervated PCs in the proximal zone of grafted cerebellar plates from explants of WT or Cry1^-/-^Cry2^-/-^ (DKO) embryos. In DKO explants, BHFS (n = 7) did not induce reinnervation compared to sham (n = 10) (1-way ANOVA with Tukey *post hoc*; p=0.946), whereas BDNF (n = 7) did (p=0.000). “WT BHFS” is given for comparison from Fig 2.

To clarify the involvement of cryptochrome in LI-rMS induced climbing fibre reinnervation, we examined the effect of its direct target, the *Clock/Arntl* transcriptional complex^24^, on gene expression. An *in silico* search of the 22 genes regulated by LI-rMS showed that 8 (36%) contained CLOCK or ARNTL binding sequences in their upstream promoter regions (Fig S4), thus supporting cryptochrome involvement in LI-rMS induced reinnervation.

## Discussion

We show that LI-rMS induced olivocerebellar reinnervation *in vivo* and *ex vivo*, similar to neurotrophin-induced repair^14,15^. Reinnervation depended on biorhythmic patterns of stimulation which upregulated expression of growth promoting genes. Reinnervation also required the presence of cryptochromes, indicating a new repair mechanism that depends on response to the magnetic field.

### Reinnervation depends on magnetic stimulation pattern

To our knowledge, this study is the first to demonstrate that LI-rTMS (*in vivo)* and LI-rMS *(ex vivo)* induce axonal outgrowth and reinnervation in a mature neural system (Figs 1&2), similar to that induced by neurotrophins^14,15,25^. Climbing fibre-Purkinje cell reinnervation requires axonal outgrowth over several millimetres, enough to have direct relevance to human neural pathology and injury. This growth contrasts with previous studies on magnetic stimulation that have only demonstrated local neurite terminal sprouting^7,26^, and suggests that magnetic stimulation has greater potential for human treatment than previously reported.

We also show the importance of stimulation frequency and pattern on post-lesion neural repair. Although human high-intensity rTMS stimulation frequencies are broadly divided into “excitatory” and “inhibitory”, with iTBS and cTBS inducing the most consistent and long-lasting effects^1,5,6,27^, our data add a new dimension to this concept. We show that complexity (iTBS vs. 10 Hz) and regularity (iTBS vs. random-iTBS) are more important than the number of pulses *per se* (random-iTBS vs. cTBS). Indeed, the same number of pulses delivered in different patterns had contrasting effects: iTBS induced Purkinje cell reinnervation whereas random-iTBS did not. These results demonstrate the critical importance of biologically-relevant stimulation patterns, such as regular runs of theta activity, in the induction of neural repair.

Although two studies have previously shown altered gene expression following magnetic stimulation^10,28,29^, this report is the first demonstration that specific stimulation patterns regulate gene expression appropriately for producing neural circuit repair. Changes in gene expression following LI-rMS are not the same for all stimulation protocols: iTBS and BHFS each stimulate the expression of genes known to contribute to neural circuit repair, especially axon growth and neural plasticity (Table S1; Fig S3). Among the functions of these gene products: β-glucuronidase (*gusb*) reduces extracellular matrix glycosaminoglycans to permit post-lesion axon outgrowth^20^, β-2-microglobulin (*b2m*) promotes axon regeneration and selective synapse stabilization^18^, and NT3 (*Ntf3*) enhances circuit repair through axon growth and synaptic plasticity in several pathways^19,30^. NT3 is also involved in olivocerebellar development and plasticity^31^ and its cerebellar injection induces climbing fibre reinnervation of denervated Purkinje cells^14^. The *gdnf* gene, whose product attracts axon growth-cones^32^, was down-regulated by random-iTBS, consistent with this pattern’s failure to induce reinnervation.

### LI-rMS requires cryptochrome magneto-receptors to induce olivocerebellar reinnervation

A key discovery of our study is the requirement for the cryptochrome for LI-rMS-induced neural circuit repair: stimulation of explants from cryptochrome double knockouts did not produce climbing fibre-Purkinje cell reinnervation. This is the first direct evidence that mammalian cryptochromes are necessary for LI-rMS-induced axon growth and neosynaptogenesis.

Cryptochrome confers light-dependent magneto-sensitivity in plants and insects^29,33^, but the light-*in*dependent magneto-sensitivity of mammalian neuronal cryptochromes had not previously been demonstrated. Cryptochromes are ubiquitously-expressed flavoproteins that undergo conformational change and generate a radical-pair in response to light or magnetic stimulation^24,34^. Radical pairs produce reactive oxygen species (ROS)^34^ and altered conformation could remove cryptochrome inhibition of *Clock/Arntl* transcriptional activity^24,35^. Because ROS control many cellular processes^36^ and CLOCK/ARNTL regulate numerous genes^37^, cryptochrome has many cellular and molecular effects. An *in silico* search for CLOCK or ARNTL binding sequences revealed their presence in promoter regions of several of the genes whose expression was modulated by LI-rMS (Fig S4), thus supporting cryptochrome involvement in LI-rMS.

In addition, ROS generation within physiological concentrations could explain how LI-rMS increases the number of c-fos positive cells^29,38^, despite the induced electric field being below action potential threshold^21^. Both ROS^39^ and LI-rMS^10^ release calcium from intracellular stores, which in turn activates c-fos. Moreover, redox equilibrium is disturbed in denervated neurons^40^, the pre-reinnervation status of our Purkinje cells and interneurons, which may account for the preferential c-fos activation in these two neuronal populations.

These data suggest a new framework for understanding the biological effects of low-intensity magnetic stimulation. Because low-intensity magnetic fields do not directly trigger neuronal firing, we argue that the magnetic field itself, acting through cryptochromes, activates a broad range of cellular events. Direct action of the magnetic field represents a shift from current thinking about high-intensity human rTMS, in which altered neuronal activity induces the plasticity. This new framework not only explains clinical^2^–4 and experimental^7,10^–13 findings, but also opens the potential to develop specific LI-rTMS programs for individual neurological conditions.

## Conclusion

LI-rTMS induces axon outgrowth and targeted neosynaptogenesis in olivocerebellar reinnervation, both *in vivo* and *ex vivo*. This reinnervation depends on cryptochrome magneto-receptors and biologically relevant stimulation patterns. Our data suggest that LI-rTMS acts on cryptochrome to trigger intracellular signalling, including changes in gene expression, that contributes to neural repair. Because cryptochrome activation can act on many intracellular targets, our findings point to the possibility of developing LI-rTMS protocols that target a range of biological questions and clinical challenges.

## Materials and Methods

### Animals

Adult C57Bl/6j mice were purchased from the Animal Resources Centre (Murdoch, Western Australia) and timed pregnant Swiss mice from Janvier-Labs (Villejuif, France). Cryptochrome knockout mice (Cry1^−/−^Cry2^−/−^, DKO) were a generous gift from Dr Xiao-mei Li and Prof Francis Lévi. Cry1^−/−^Cry2^−/−^ embryos were obtained from timed mating between Cry1^−/−^Cry2^-/−^ male and Cry1^−/−^Cry2^+/−^ females. The mouse phenotype has been described previously^41^.

Briefly, tissue was digested overnight at 55°C with proteinase K (Qiagen, France) in TSE buffer containing: 25 mM Tris–HCl pH 8.0, 75 mM NaCl, 25 mM EDTA pH 8.0, and 1% SDS (Sigma-Aldrich, France). DNA fragments were precipitated with isopropanol and washed with 70% ethanol prior to being dissolved in 100 µl of distilled water. DNA was amplified by PCR and the amplified fragments were detected by electrophoresis on a 2% agarose gel. The following primers were used:

Cry1 P1 (wt) Ex5: 5’-TGA GGC ACT TAC ACG TTT GG-3’

Cry1 P2 (ko): 5’-TGA ATG AAC TGC AGG ACG AG-3’

Cry1 P3 wt/ko: 5’-ATC CCT TCT TCC CAG CTG AT-3’

Cry2 P1 (wt): 5’-CCA GAG ACG GGA AAT GTT CTT-3’

Cry2 P2 (ko): 5’-GAG ATC AGC AGC CTC TGT TCC-3’

Cry2 P3 (wt/ko): 5’-GCT TCA TCC ACA TCG GTA ACT C-3’

Animal housing and all procedures were authorized either by the Comité National d’Ethique pour les Sciences de la Vie et de la Santé (N° 1492-02) or the University of Western Australia Animal Ethics Committee (N° 03/100/834), in accordance with the European Communities Council Directive 2010/63/EU and regulations of the NH&MRC of Australia and the NIH.

### In vivo olivocerebellar axonal transection (pedunculotomy)

Our repair readout is the post-lesion restoration of climbing fibres, which connect the inferior olivary nucleus to Purkinje cells of the cerebellum^14–16^. Unilateral climbing fibre transection leads to degeneration of the axotomized inferior olive. Intra-cerebellar injection of neurotrophic factors can induce neurons of the remaining inferior olive to develop axon collaterals that partially reinnervate the denervated hemicerebellum and compensate motor coordination and spatial learning behaviours^14,15^.

WT mice were anaesthetized with Xylazine (10 mg/kg i.p.) and Ketamine (70 mg/kg i.p: Ilium, New South Wales Australia) and underwent unilateral transection of the left inferior cerebellar peduncle as previously described^14,15^. Briefly, the skin over the neck was incised longitudinally and the muscles retracted to expose the atlanto-occipital membrane. A capsulotomy knife (MSP, 3mm blade) was inserted parallel to the brainstem into the fourth ventricle and rotated to the left to cut the left inferior cerebellar peduncle. After recovery from the anaesthetic, animals were returned to the cage. Food and water were provided ad libitum.

### Organotypic cultures and cerebellar denervation

The olivocerebellar pathway can also be studied ex vivo using organotypic hindbrain explants that contain the whole circuit, are highly reproducible and readily manipulated^17,42^, so that we can reproduce the pedunculotomy lesion. Hindbrain explants were cultured from embryonic Swiss or Cryptochrome knock-out (Cry1^-/-^Cry2^-/-^) mice at embryonic day 14 (E14) as previously described^17,42^. E0 was the mating day. Following anaesthesia and cervical dislocation of pregnant females, embryos were removed and their brains quickly dissected in ice-cold Gey’s balanced salt solution (Eurobio) containing 5 mg/mL glucose. The hindbrain, including the cerebellar anlage and inferior olivary nucleus, was isolated and the meninges removed. The right and left cerebella plates were separated at the midline (Fig S1) and the explants transferred onto 30mm Millicell membranes (pore size 0.4µm, Millipore) then cultured with medium, containing 50% basal medium with Earle’s salts (Gibco), 2.5% Hank’s Balance Salt Solution (Gibco), 25% horse serum (Gibco), 1mM L-glutamine (Gibco), and 5mg/mL glucose, at 35°C in humidified air with 5% CO2. The culture day was designated 0 day *in vitro* (DIV). The medium was replaced every 2–3 days.

To denervate cerebellar tissue and induce olivocerebellar reinnervation, we removed the cerebella plates from their explant brainstem at 21 DIV (equivalent to P15) and co-cultured them (graft) adjacent to the cerebellar plate of an intact explant (host; Fig S1).

### Magnetic stimulation

#### In vivo

Pedunculotomised mice recovered for 3 days post-surgery before receiving LI-rTMS or sham treatment, 10 minutes per day for 14 consecutive days, as previously described^8,9^. An electromagnetic pulse generator (EC10701, Global Energy Medicine, Perth, Australia) was modified for attachment of a custom-designed copper wire (0.25 mm diametre, 16Ω; Jaycar) coil with an outer diameter of 8mm, 300 windings and a 6mm diameter steel bolt in the centre to increase field penetration^9^. The non-sinusoidal monophasic 300µs pulse had a measured 230µs rise time and generated a magnetic field intensity of 12 mT at its surface, without sound, vibration or heat above background^9,38^. The coil size was designed to ensure a similar coil-to brain ratio as used for focal magnetic stimulation in humans^43^.

#### Ex vivo

To investigate which parameters were most effective for inducing reinnervation, pedunculotomised explants received LI-rMS (identical to LI-rTMS but not "transcranial") inside the incubator, 10 min per day for 14 days, using a custom-built copper wire coil (10 mm inside diameter, 26 mm outside diameter, 199 turns) placed 4 mm below the well and driven by a 24 V magnetic pulse generator^38^. The non-sinusoidal monophasic 300µs pulse had a measured 100µs rise time and generated an intensity of 10 mT at the explant without sound, vibration or heating above background^38^. Each culture plate was isolated using mu-metal (nickel/iron/molybdenum alloy) shielding to ensure no eddy current spill over. In some experiments mu-metal was used to shield either the rostral or caudal part of the explant from the stimulation.

Electric and magnetic fields were modelled using the finite element package MagNetv7.0 (Infolytica, Inc., Canada) as previously described^38,44^. As previously demonstrated the modelled electric field in the explant’s regions of interest (cerebellar lobes and inferior olive nucleus) was 0.05 Vm^-1^. These electric fields are at least two orders of magnitude below the electric field amplitude reported for activation of cerebellar neurons^21^.

We chose stimulation patterns (Fig S1) that are used in human clinical rTMS: simple pulse stimulation at 1 Hz and 10 Hz; complex frequencies with a theta-burst stimulation pattern (TBS; 3-pulse bursts at 50 Hz repeated at 5 Hz), delivered either continuously (cTBS), intermittently (iTBS) for 2 s repeated every 10 s, or randomly (random-iTBS: 3-pulse bursts at 50 Hz repeated randomly at 2-60 Hz, for 2 s, all repeated every 10 s). We also tested a complex biomimetic high-frequency stimulation (BHFS), which we have previously shown can modulate neural circuits^8,9^. Sham treatment was delivered identically but without activation of the stimulation coil.

### Immunohistochemistry

Pedunculotomized mice were euthanized 24h after the last stimulation (18d post-lesion) with an overdose of sodium pentothal and perfused transcardially with 0.9% saline and 4% paraformaldehyde in 0.1M phosphate buffer. The brain was post-fixed in fresh 4% paraformaldehyde overnight at 4°C, cryoprotected in 30% sucrose, and coronal cryosections of the cerebellum and brainstem cut at 20 µm. For ex vivo experiments, 24h after the last stimulation session explants were fixed with 4% paraformaldehyde for 4h at 4°C.

Sections and explants were labelled by immunohistochemistry. Fixed tissue was rinsed 3×5 min in phosphate buffered saline (PBS) containing 0.25% Triton X (PBS-T) and blocked in 20% donkey serum for 2h at RT prior to incubation overnight at 4°C in primary antibody diluted in PBS-TG (PBS-T containing 0.2% gelatine and 0.018g/mL L-Lysine). The next day sections or explants were washed 3×5 min in PBS-T and labelling was visualised with fluorescent-conjugated secondary antibodies in PBS-TG for 2h at RT. Finally, sections or explants were rinsed and mounted in Mowiol.

To identify climbing fibre reinnervation, Purkinje cells were labelled with rabbit anti-calbindin-28k (CB) antibody (1:3000; Swant) and climbing fibre terminals with polyclonal guinea pig anti-VGLUT2 antibody (1:2000; Millipore)^14,15,17^. Primary antibodies were visualised using Cy3-conjugated donkey anti-guinea pig and Alexa Fluor (AF) 488-conjugated donkey anti-rabbit (1:200 and 1:400 respectively; Jackson Laboratories).

To identify which cells were activated by the magnetic stimulation ex vivo we labelled for cfos 2.5h after a single 10 min stimulation session 72h post-lesion and co-culture. Fixed explants were labelled with rabbit anti c-fos (Synaptic System, 1:3000) and one of 4 different antibodies to identify specific cell populations^45^: Purkinje cells with monoclonal mouse anti-CB (1:2000; Swant), GABAergic interneurons with goat anti-parvalbumin (PV, 1:3000; Swant), granule cells with monoclonal mouse anti-NeuN (1:200; Millipore). Primary antibody binding was visualised using Cy3-conjugated donkey anti-rabbit, AF488 conjugated donkey anti-goat and Cy5 conjugated donkey anti-mouse (all 1:200; Jackson Laboratories).

### Histological Analysis

#### Quantification of olivocerebellar reinnervation

In vivo experiments: Immuno-labelled sections were examined using epifluorescence microscopy (E800; Nikon). To quantify the extent of reinnervation, the cerebellum of pedunculotomized animals was divided into a series of parasagittal zones (500 µm wide) extending from the midline to the left lateral hemicerebellum. Within each zone, the amount of VGLUT2-positive climbing fibre reinnervation was scored in each lobule using an arbitrary scale, i.e. 1 = few strands, 2 = one-quarter climbing fibre-filled lobule, 3 = half lobule, 4 = three-quarters lobule and 5 = completely climbing fibre-filled lobule. Scores in each 500 µm zone of each lobule are the mean value from all animals and are plotted on an unfolded cerebellar cortex to give a semi-quantitative evaluation of climbing fibre reinnervation as described previously^15^.

Ex vivo experiments: Labelled explants were examined using epifluorescence microscopy (DM 6000F; Leica) and z-stack images taken for analysis. Climbing fibre reinnervation was quantified by counting the number of CB-positive Purkinje cells (soma and primary dendrites) co-localised with VGLUT2 per field of view, and expressed as percentage Purkinje cells per field. This quantification was made systematically on z-stacks taken in rows through the cerebellar graft with increasing distance from the host-graft interface (Suppl Fig 1). Data from rows 1 and 2 were defined as the proximal zone, and those from rows 3-5 were defined as the distal zone.

#### Cellular activation: c-fos

Explants were examined using epifluorescence microscopy and z-stack images were taken at 3 systematic randomly-selected sites of 0.073 mm2 for each co-cultured cerebellar plate. Total c-fos positive staining was counted per z-stack and double-labelled or triple-labelled profiles were examined for co-labelling. C-fos, CB and PV positive cells were counted per image to identify the proportions activated by LI-rMS. Results are expressed as mean number of c-fos positive cells per mm2.

### qRT-PCR

Changes in gene expression, triggered by different LI-rMS frequencies, were examined in a separate series of explants following effective and ineffective reinnervation protocols. Explants were denervated and co-cultured at 21 DIV (P15) and 72h after the lesion they received 3 daily sessions (10 min/day) of LI-rMS/sham. The cerebellar plate of denervated-treated explants was taken 6h after the last stimulation (P18). Six cerebellar plates were pooled and total RNA was extracted using Trizol (Life Technologies) according to manufacturer’s instructions and stored at −80°C.

cDNA was transcribed from 400 ng total RNA using the RT² Easy First Strand cDNA Synthesis Kit (Qiagen). For each sample, the resulting cDNA was applied to the RT² Profiler™ PCR Array Mouse Neurotrophins & Receptors (Qiagen) and amplified with a Lightcycler480 System. Results were analysed on the Qiagen RT2 Profiler PCR array data analysis (v3.5) using the housekeeping gene, hsp90ab1. Normalized mean expression levels (log2(2−ΔCt)) were used to determine differentially expressed genes between each group and control.

### Statistical Analysis

All data were explored for normality, outliers and fulfilment of statistical test assumptions in SPSS 22 (IBM Corp, Armonk, NY). Reinnervation percentages were analysed with Repeated Measures ANOVA (F) (location x stimulation). When significant interaction between stimulation and location were present, univariate ANOVA were performed to evaluate the effect of the treatment in proximal and distal zones independently. Tukey post-hoc comparisons were performed where appropriate.

C-fos activation was analysed with univariate ANOVA. Tukey post-hoc comparisons were performed where appropriate. CB/c-fos PV/c-fos were analysed with Fisher exact test. Gene expression levels were compared by two-sample t-test. All values are expressed as mean ± SEM and considered significant at p < 0.05.

The function and linkage of regulated genes were examined for GO terms enrichment using the R topGo package. Searching for GO biological process, molecular function or cellular component (GO.db 3.3) each list of genes was analyzed by the topGo over representation test against the Mus musculus reference list. The Bonferroni correction for multiple testing was applied. Using R (3.3.0) and the Bioconductor suite (3.3), we applied Biostrings (2.40) and the GenomicFeatures (1.24) to determine potential transcription factor (TF) binding sequences in our regulated genes. TF binding matrices (PWM) were obtained from the R package MotifDb (1.14) and compared, using matchPWM (with Position at 90%), to target sequences in the promoter site and for 2 kb upstream

## Acknowledgments

We thank Prof Allan Tobin for his help with the manuscript.

## Funding

This work was supported by the Institut pour la Recherche sur la Moelle Epinière et l’Encéphale, the Centre National de la Recherche Scientifique (CNRS) and the Paris region Ile de France (CPER). T. Dufor and S. Grehl had PhD scholarships from the French Ministry of Education and the University of Western Australia, respectively. Z.-D. Deng is supported by the National Institute of Mental Health Intramural Research Program, NIH;

## Author contributions

RMS and TD designed the experiments; TD, SG, AT and RMS did the experiments; MD, ND and ZDD undertook bioinformatics and electromagnetic field modelling; TD, MT, CD and MD did the molecular biology; TD, ZDD, JM, AML and RMS analysed results and wrote the paper;

## Competing interests

Authors declare no competing interests.;

## Data and materials availability

All data are available in the main text or the supplementary materials.

## Supplementary Materials

Figures S1-S4

Tables S1, S2

## Supplementary Materials for

**This PDF file includes:**

Figs. S1 to S4

Tables S1 and S2

**Fig. S1:**
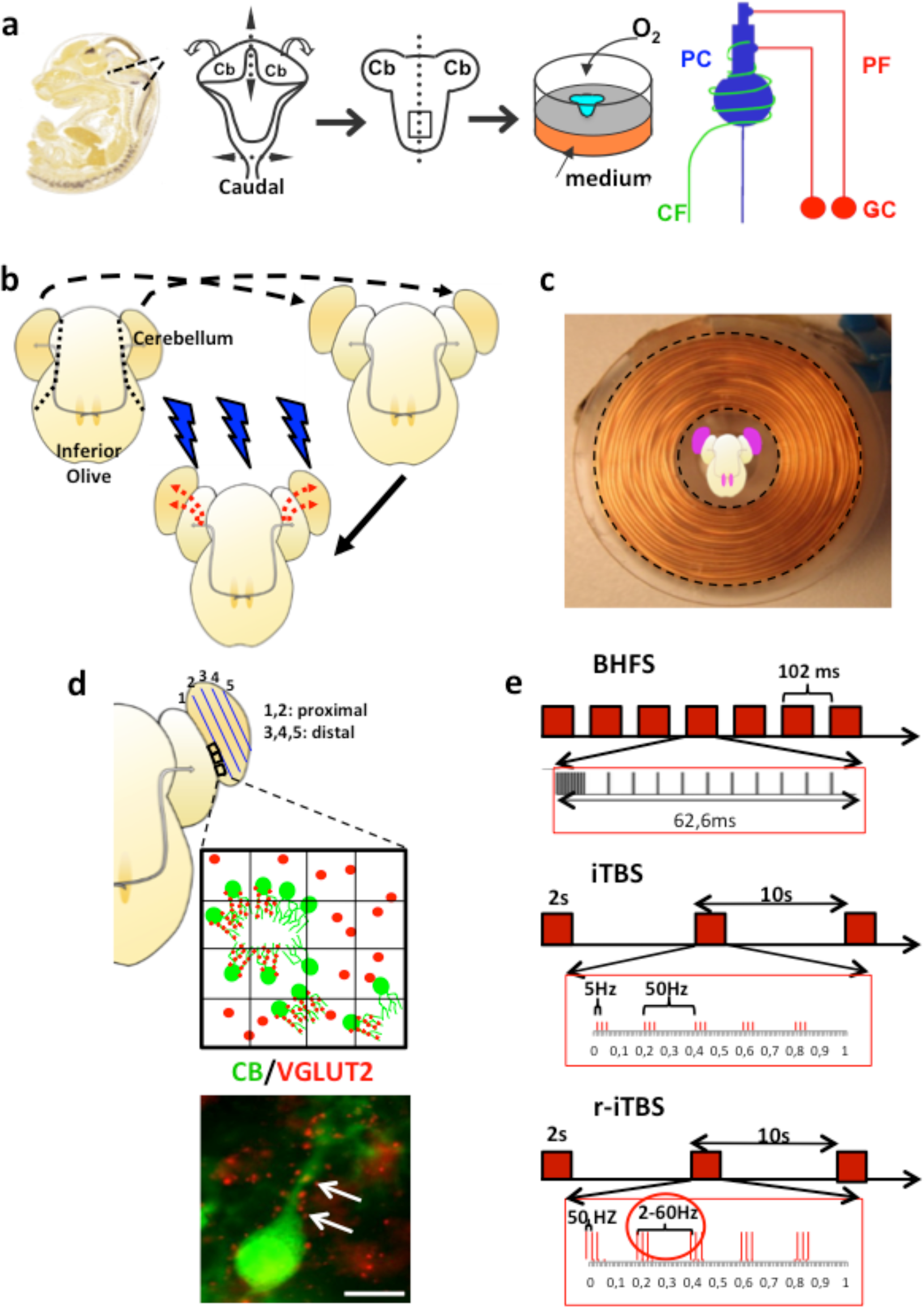
Experimental protocol for LI-rMS to an *ex vivo* model of the olivocerebellar path. **a.** Diagrams of the dissection and culture of embryonic mouse hindbrain. The hindbrain from an E14 mouse embryo is quickly dissected and the meninges removed. The right and left cerebella plates were separated at the midline and the explants transferred onto 30mm Millicell membranes then cultured at 35°C in humidified air with 5 % CO2. These explants develop the cerebellar cortical circuit as *in vivo*, so that Purkinje cells develop branched dendritic arbors that receive parallel and climbing fire afferents ^1,2^. **b.** Diagram of the pedunculotomy and co-culture procedure. Hemicerebellar plates are removed from an explant (black dotted line) at 21div (equivalent to P15) and placed next to the hemicerebellar plates of another intact explant (black dashed arrows). LI-rMS-induced collateral sprouting of climbing fibre axons (thin grey arrows) into the denervated hemicerebellar plate (red dotted arrows) can then be studied. LI-rMS = 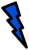 **c.** Schematic representation of the spatial relation between the LI-rMS stimulation coil and the olivocerebellar explant. The lesioned (denervated) hemicerebellar plates and the ION (both in purple) are stimulated by a homogeneous magnetic field (as previously described; ^3^). **d**. Schematic diagram of climbing fibre reinnervation analysis. Hemicerebellar plates were divided into 5 rows (1-5) parallel from the host-graft junction. Within each row, PCs (CB staining in green) found within an ocular grid field are checked for CF reinnervation (small VGLUT2-positive puncta (red, indicated by the white arrows in lower image). VGLUT2-positive mossy fibres are larger red-labelled terminals not localised on the PC. Data from rows 1 and 2 are pooled and designated the proximal region, and data from rows 3, 4 and 5 were pooled and designated the distal region. The white scale bar is 20µm. **e.** Diagram of the different “complex” patterns of LI-rMS tested in this study. BHFS delivers trains of 20 pulses lasting 62.6ms (the first 9 pulses at 1.4kHz and the last 11 at 188Hz) repeated at a 9.75Hz^4,5^. iTBS consist of bursts of three pulses at 50Hz repeated at 5Hz. Trains of bursts last 2s and are repeated every 10s. cTBS is similar to iTBS but the trains of burst are delivered continuously for the 10mn of stimulation. Random-iTBS (r-iTBS) is similar to iTBS but the bursts are repeated at random frequencies ranging from 2-60Hz.

**Fig. S2:**
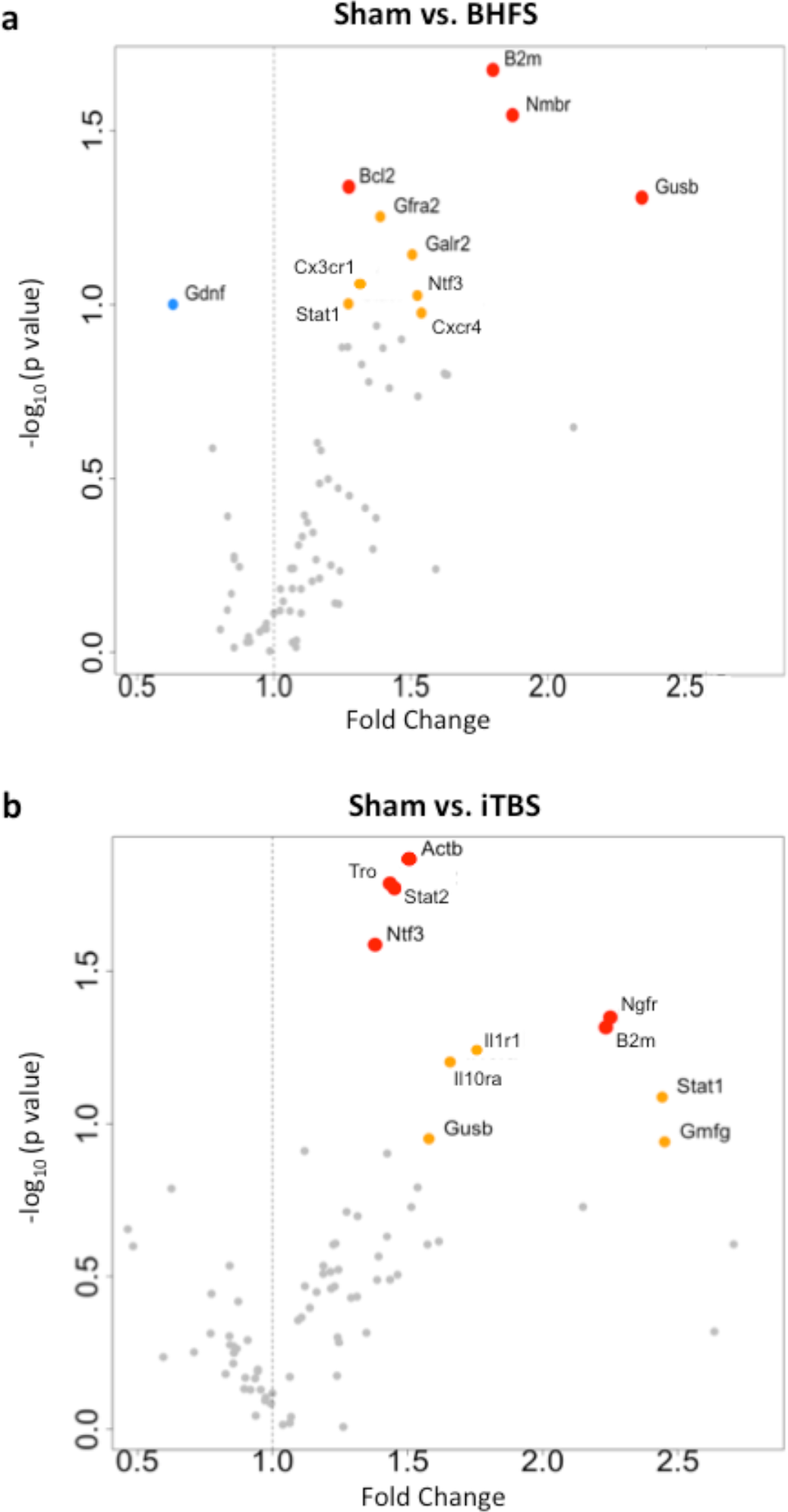
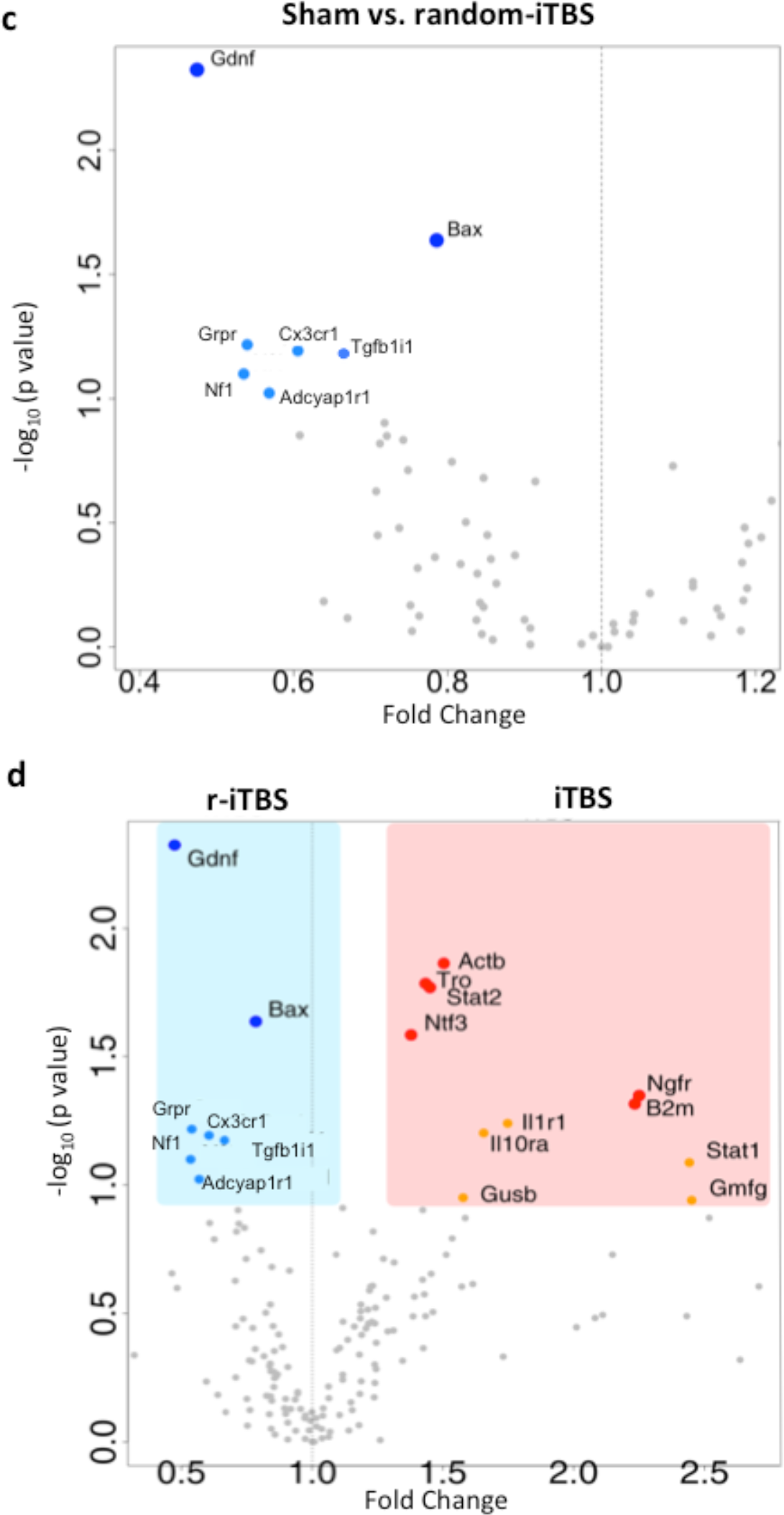
Specific LI-rMS patterns modulate gene expression appropriately for PC reinnervation. Volcano plots showing changes in expression of genes from the Mouse Neurotrophins & Receptors PCR array (Qiagen), that were induced by BHFS (a), iTBS (b) and random-iTBS (c) compared to sham treatment (*n*=5 for each group). (d) is a summary to show the clear difference between the gene changes induces by iTBS (pink shading) and random-iTBS (pale blue shading). Red dots represent genes that are significantly upregulated (p<0.05); orange dots represent genes that show a strong trend for upregulation (0.05<p<0.1); royal blue dots represent genes that are significantly down regulated (p<0.05); light blue dots represent genes that show a strong trend for down regulation (0.05<p<0.1). Small grey dots represent genes that are not strongly regulated by LI-rMS.

**Fig. S3:**
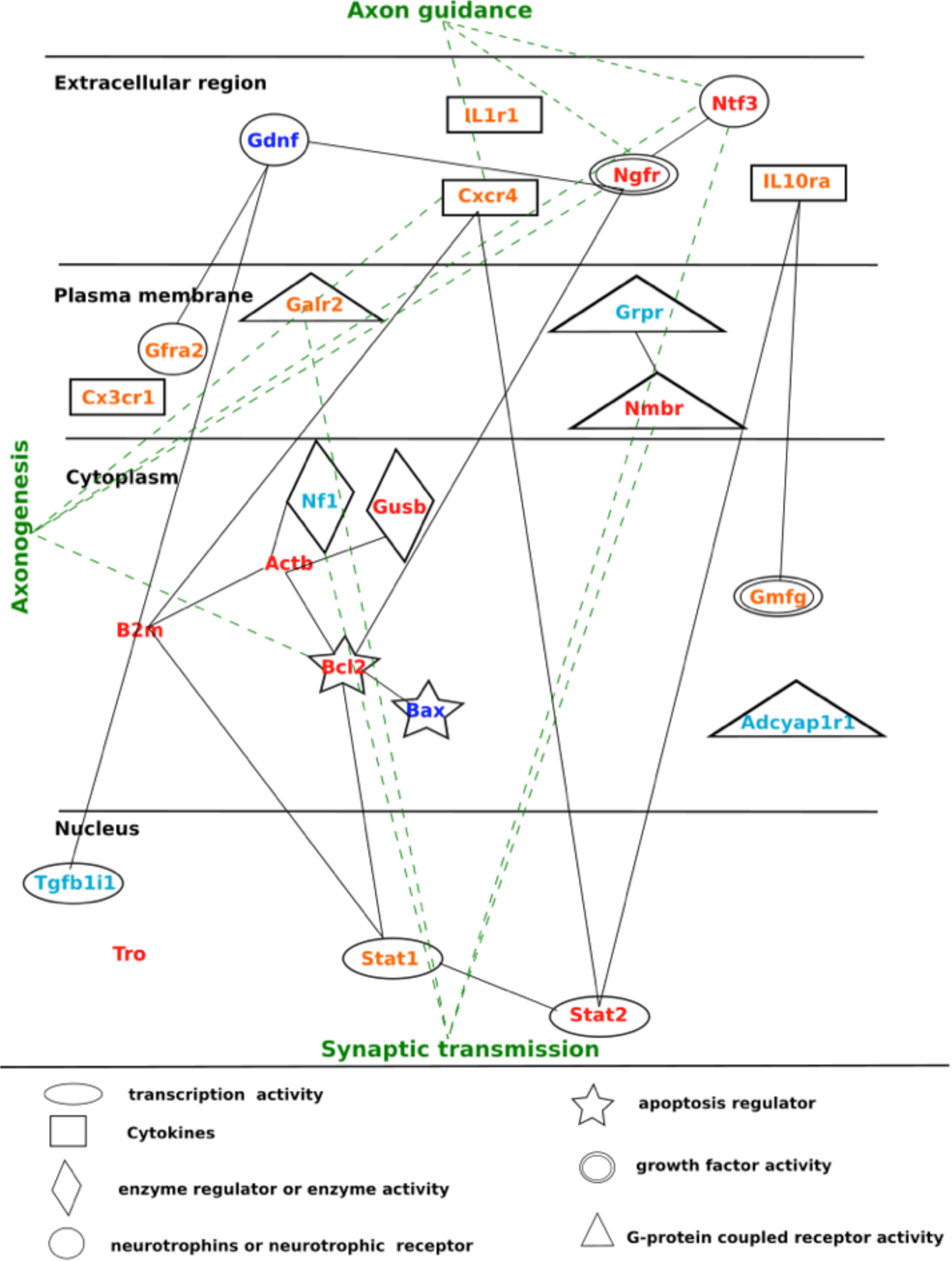
Relationships between genes regulated by different LI-rMS patterns. Pathway diagram obtained from STRING showing the relationships between genes regulated by different LI-rMS patterns in our study and the location of their function. Also, genes are linked to their main GO function by green dotted lines. As indicated in Table S1 several genes are involved in more than one GO group. The colour code of gene names represents the expression changes as described in Figures 4, S2 and Table S1: Red represents genes that are significantly upregulated (p<0.05); orange genes that show a strong trend for upregulation (0.05<p<0.1); royal blue represents genes that are significantly down regulated (p<0.05); light blue represent genes that show a strong trend for down regulation (0.05<p<0.1).

**Fig. S4:**
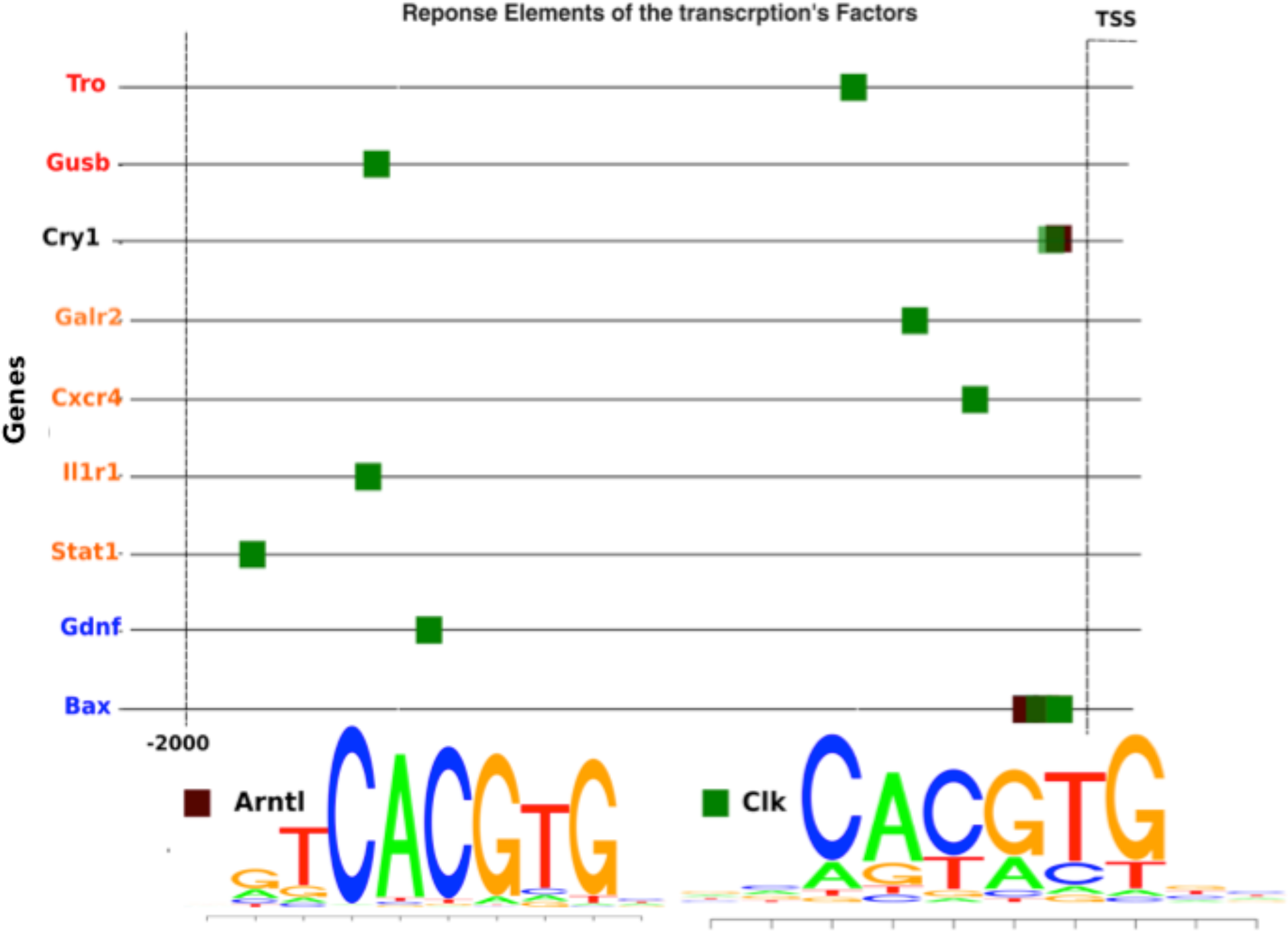
Genes regulated by different LI-rMS patterns contain binding sites for the transcription factors CLOCK and Arntl1. Predicted transcription factor (TF) binding sites for CLOCK and Arntl1, which are direct targets of Cry, were searched in the DNA sequences up to 2kbp upstream of the transcription starter site (TSS) for each gene regulated by LI-rMS. The position weight matrices for Arntl1 and CLOCK indicate the presence of specific binding sites for either of these two TFs in the upstream promoter region, and the probability of specific bases at each position within the TF response elements.

**Table S1.**
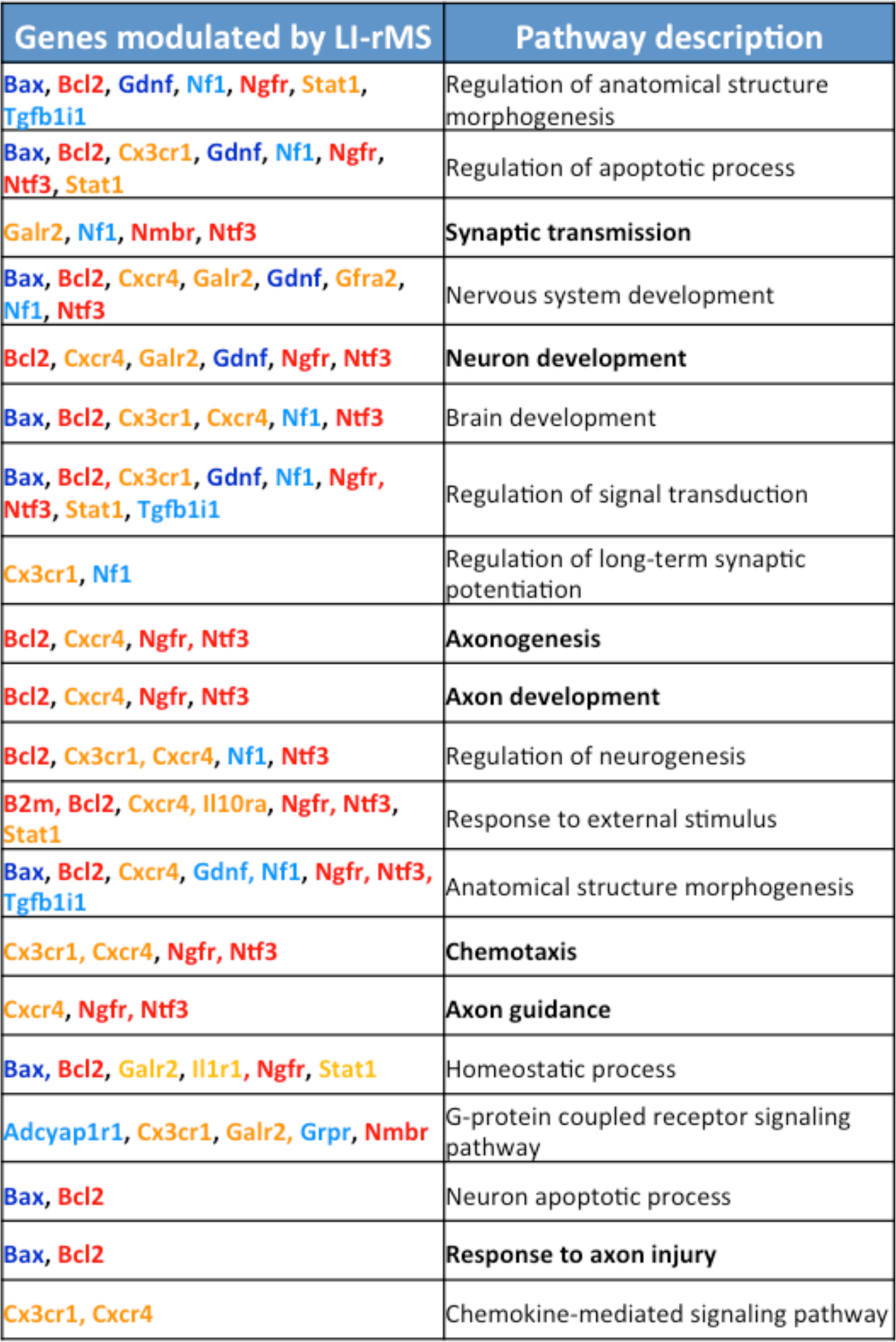
Biological pathways of genes regulated by LI-rMS. Biological pathways in which LI-rMS regulated genes are implicated. The colour code is the same as in Figures 4d&e, S2 and S3. Gene ontology (GO) terms enrichment was assessed using an *in silico* approach against the *Mus musculus* reference list, using GO biological process, molecular function or cellular component. The Bonferroni correction for multiple testing was applied.

**Table S2.**
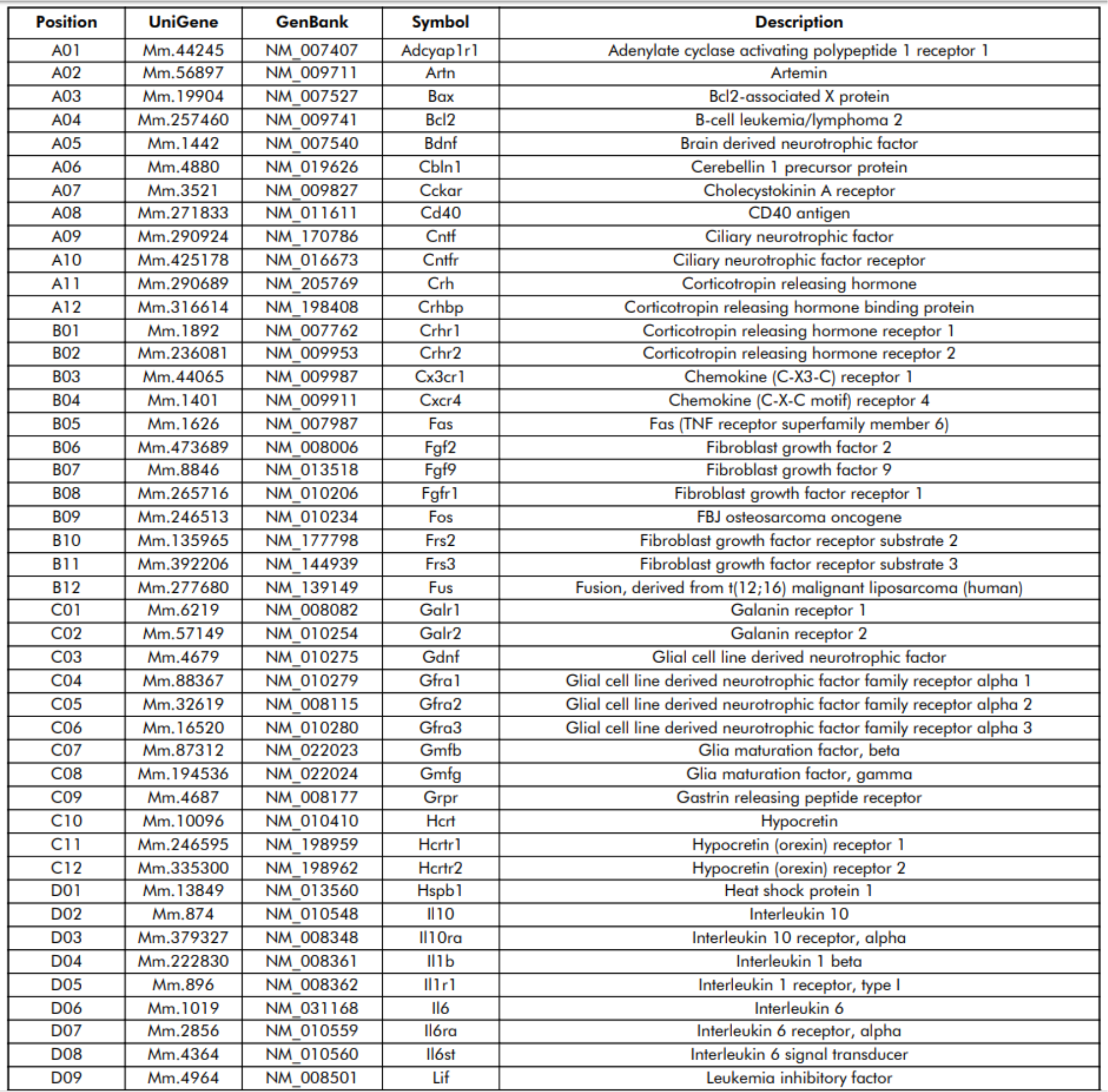
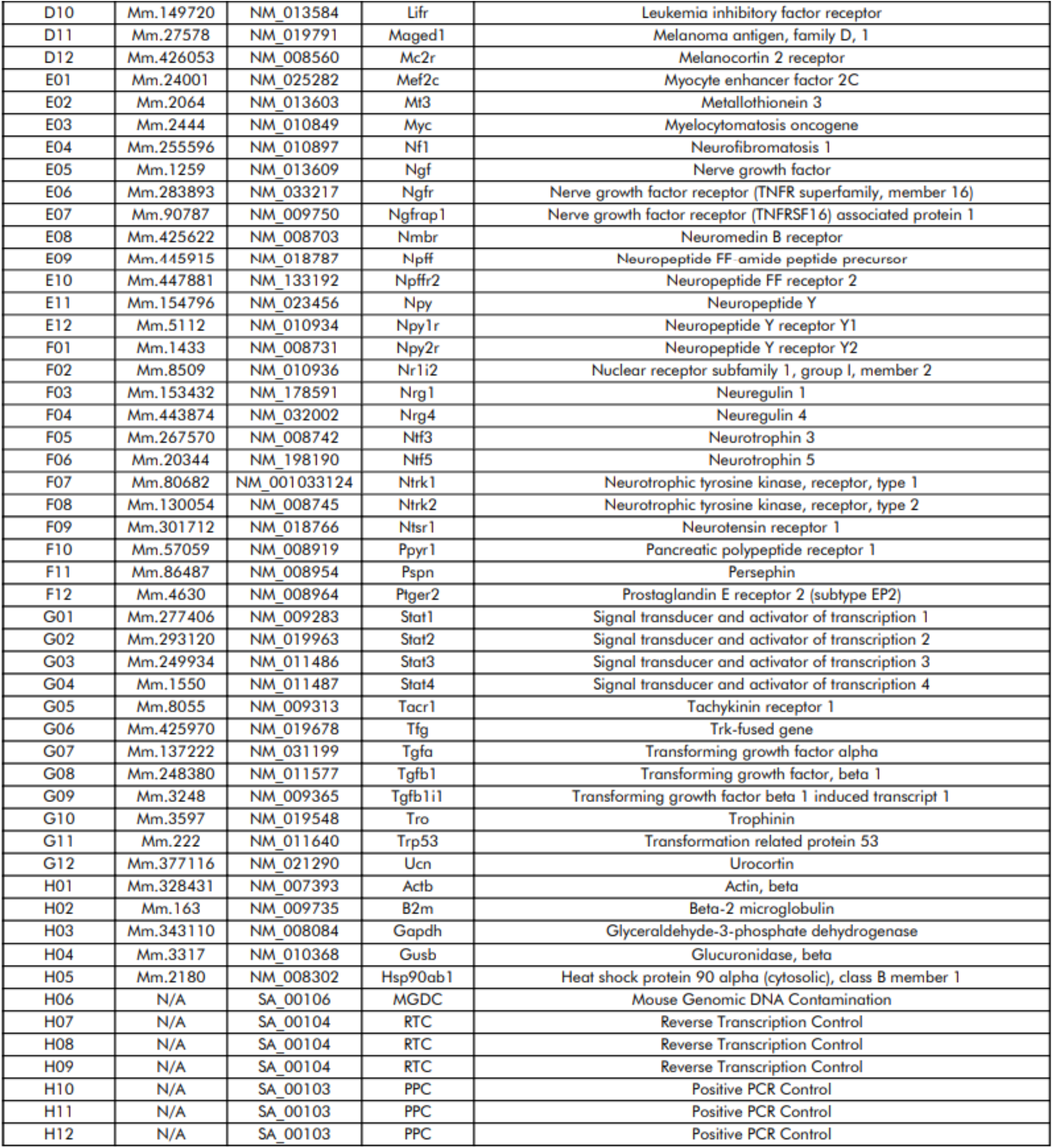
Genes examined on the RT² Profiler™ PCR Array Mouse Neurotrophins & Receptors.

